# Arrest of movement induced by Pedunculopontine-stimulation obstructs hippocampal theta rhythm

**DOI:** 10.1101/2023.08.26.554923

**Authors:** Jaspreet Kaur, Salif A. Komi, Oksana Dmytriyeva, Grace A. Houser, Madelaine C. A. Bonfils, Rune W. Berg

## Abstract

Complete movement arrest has recently been reported to be induced in rodents by optogenetic stimulation of a subpopulation within the pedunculopontine nucleus (PPN). This evoked arrest appears conspicuously similar to freezing behavior often seen as a fear response in prey animals but could also be akin to the freezing of gait, which is a symptom of Parkinson’s disease. This introduces two perspectives on the functional roles of this sub-region: A hub for orchestrating fear-related responses or an omnipotent halting mechanism devoid of emotional components. To better understand this phenomenon and its cognitive component, we engage the distinct electrical brain activity, the hippocampal theta rhythm. This rhythm has a well-described contextual association between various aspects of cognition and behavior. It is prominent during locomotor activity in rodents and immobile yet aroused states like behavioral freezing. We recorded the electrical activity in the hippocampus of rats while walking and being arrested by PPN stimulation. A clear obstruction of the ongoing theta activity was associated with the motor arrest. The timescale of movement arrest was less than 200 ms, similar to the obstruction of the theta rhythm. Since, anxiety, fear, and behavioral freezing are associated with hippocampal theta rhythm, which we did not see during PPN stimulation, we suggest the induced motor arrest occurs without an associated emotional component.

---

A temporary arrest of movement is a natural part of animal and human behavior^1^. For instance, an unexpected event or the appearance of an obstacle would require suppression of all movement^2^. It is also seen for example, in rodents as a freezing behavior to a pain-conditioned response or looming threat^3^ with an immediate need for silence and immobility. It was recently observed that a similar arrest of movement could be induced by optogenetic stimulation of a subset of glutamatergic neurons, the Chx10 cells^4,5^, in a nucleus in the caudal mesencephalon known as the PedunculoPontine Nucleus (PPN). Similar behavior was observed by stimulation of unidentified neurons in PPN both in mice^6^ and rats^7^ and reduced motor activity and long-lasting muscle tone in mice^8^ and cats^9,10^. These observations were remarkable since, the PPN is part of a functionally defined region, which is known for inducing locomotion, not suppression of movement. This is the midbrain locomotor region (MLR), which was first identified in cats in an early study^11^. By means of electrical stimulation, the investigators could consistently evoke locomotion as long as the stimulation pulse train remained. An increase in stimulus intensity induced faster locomotion in terms of stepping frequency and type of gait. The PPN together with another brainstem nucleus, the cuneiform (CnF), comprise the MLR. The CnF and PPN have similar properties regarding evoking locomotion albeit with a discrepancy in slow exploration versus high-speed escape locomotion^6,12^. The MLR has descending fibers that are relayed in the reticular formation that further descend to activate rotational population dynamics in spinal central pattern generators responsible for the production of movement^13^. The capacity of MLR to activate the central pattern generation has been extensively confirmed in various species and experimental conditions^14–22^. PPN also has a central role in orchestrating behavioral states in environmental contingencies^23–25^ as well as sympathetic cardiovascular control^26^. Thus, the observation of a global movement arrest induced by optogenetic stimulation in PPN was unexpected. Due to the connection between PPN and the basal ganglia^8,27–29^, the resemblance of the evoked response with the Parkinsonian symptom of bradykinesia and freezing of gait is also clinically interesting^30^. The MLR has been suggested to have a pathophysiological role in Parkinson’s disease and a therapeutic target^31–33^. The discrepancy of reports of the evoked responses to PPN stimulation has been demonstrated to be related to both the location within the PPN as well as the cell type being activated^5,7,10,17^. Nevertheless, the movement arrest by stimulation of the PPN raises new questions on the functional significance of this nucleus. Is the evoked movement arrest tied to a feeling of startle or fear? If not, what is the cognitive state during the arrest? Movement arrest is seen when activating neurons in the central amygdala^34,35^ and thus an emotional coupling to the PPN-evoked movement arrest is possible.

Movement and the location of the body in space are closely tied to the representation of events in time. Hence, locomotion has to be tightly integrated into the perception of events^36,37^. Activating PPN affects the state in the cortex^23,38^ and the hippocampus^39–41^, but it is unknown how the internally-induced movement arrest by stimulating PPN affects the processing of time and spatial location of the body. A part of the brain, which is known to have a central role in both memories of events and spatial navigation is the hippocampus^36,42^. A prominent feature of the hippocampus is the strong concerted rhythmic activity, known as the theta rhythm (6-9 Hz in rats), which is prominent, especially during movement, locomotion^43–45^ as well as the acceleration of the body^46^. It often appears together with a widespread desynchronization of cortical electroencephalography (EEG) and was therefore suggested early on to be involved in arousal^39^. It was also observed in early reports that the theta rhythm is often associated with motor activity and hence proposed to have a relationship with sensorimotor processing^47^. Nevertheless, these events, which exert modulatory effects on the hippocampal theta rhythm either as sensory- or motor-related contingencies, have so far been related to external events. The recently reported movement arrest by internal modulation of the PPN could represent a temporary “highjacking” of the motor control, which would be reflected in a continuation of hippocampal theta oscillation during the induced immobility. Since theta rhythm is present during immobile preparedness^48^ and associated with aroused and fearful states during freezing^49^, anxiety and “panic”^50^ or due to a nearby predator, e.g. a rat having visual contact with the movement of a cat^51,52^. The resemblance between the PPN-induced movement arrest and a true fear-induced freezing behavior would further strengthen the prediction of theta rhythm being present during PPN stimulation. This type of theta rhythm is also known as type-2 theta or “atropine sensitive” theta^45,53^. The alert immobility is not obviously different from the movement arrest induced by optogenetic stimulation in the PPN.

Here, we explore this internally induced arrest of movement to investigate the relationship between this brainstem area, the hippocampal activity, and the functional machinery in the motor system. We directly test the impact of the PPN-induced movement arrest on the ongoing theta rhythm during the locomotion and exploratory movement of rats.

## Results

### The arrest of movement by PPN stimulation

Our first objective was to reproduce the previously reported movement arrest by optogenetic stimulation in the PPN^4–7^. The opsin was expressed using an AAV virus injected into the PPN^7^ along with the implantation of an optical fiber (**Fig. 1A**). After expression, the animal was allowed to perform exploratory movements, which were recorded using accelerometers. When optogenetically activating the PPN the animal rapidly came to a halt (**Fig. 1B**). This is seen as the extinction of the accelerometer measurement coinciding with the PPN stimulation (“PPN stim”, blue-shaded regions). When stimulating PPN, the rat would keep its posture at the stimulus onset, which could be seen e.g. while locomoting on a treadmill (“flexed paw pose”, **Fig. 1C**). When the stimulation was stopped, the animal would resume the locomotion (**Supplementary movie 1**). This is consistent with the previously reported “pause-and-play” pattern observed in mice^5^. The arrest was quantified in terms of the accelerometer measurement in x, y, and z-directions, which gives an upper limit on the effects on the neural processes. These were triggered at the onset of the stimulus, rectified, and averaged across directions (**Fig. 1D**). For comparison, the averaged values were integrated at 300 ms after the start of the stimulus (“PPN-stimulation”) as well as at 200 ms prior to onset, which served as pre-values (“control”). The movement versus PPN stimulation was compared across stimulus events for the same animal (**Fig. 1E**) and their median value was compared across animals (**Fig. 1F, Supplementary Fig. 1**). They all show a significant reduction in movement, in terms of a decrease in accelerometer measurement when the PPN was stimulated.

**Fig. 1.**
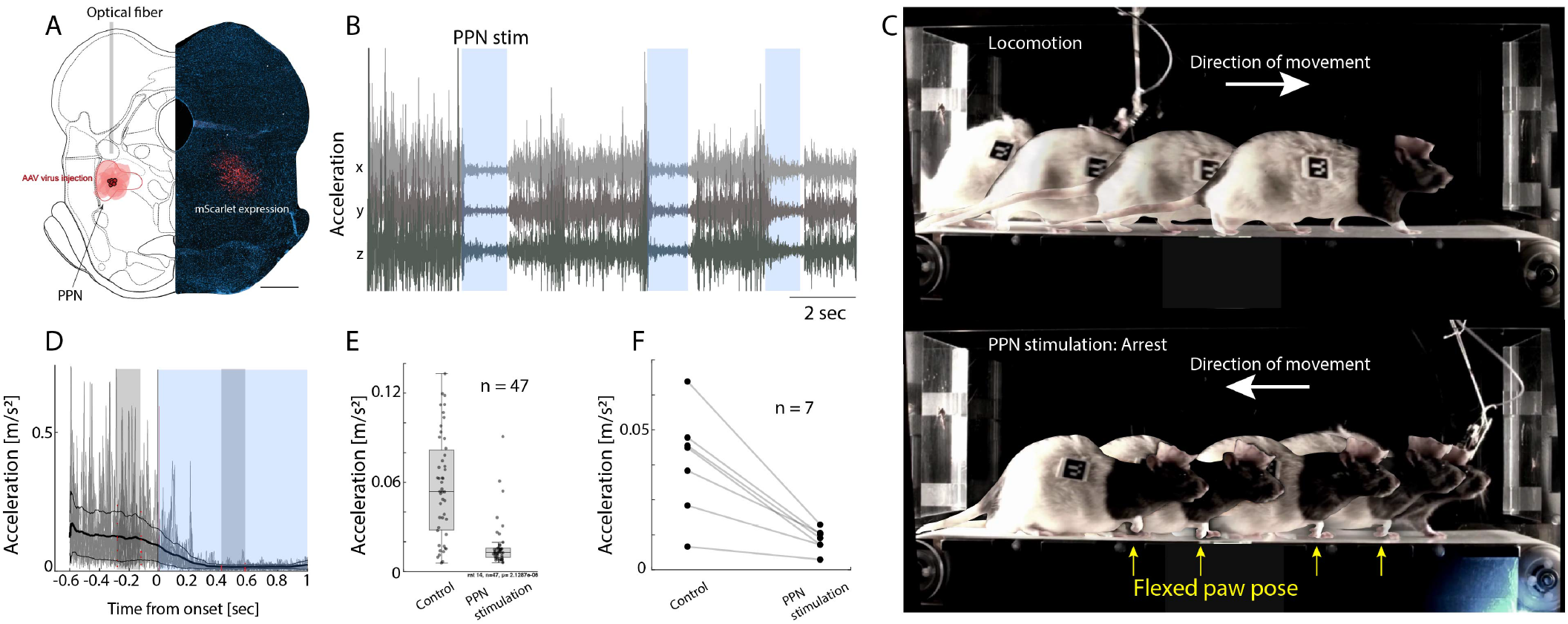
Optogenetic stimulation of PPN arrests ongoing movement. (A) The PPN was targeted in rats using optogenetics via a virus, which expresses an opsin (ChrimsonR) and an implanted optical fiber. A histology section (scale bar= 1000 *μ*m) of the caudal mesencephalon shows the viral reporter (mScarlet) in red and DAPI staining in blue and the corresponding atlas section with injection sites in red. (B). Accelerometer readings of movement in 3 directions (anterior-posterior, mediolateral, and dorsoventral), and optical stimulation (“PPN stim”, blue regions) induced arrest and eliminated acceleration. (C) Chronophotography of rat locomotion on a treadmill. Top: While not stimulating PPN, the rat is moving faster than the belt speed, hence advancing forward. When optically stimulating the PPN, the locomotion is halted (bottom) while keeping its pose). Flexed paw pose is indicated (yellow arrows) while moving backward due to belt movement. (D) The stimulation-triggered and averaged rectified accelerometry (black line). The average is integrated across the shaded gray regions across trials as “control” and “PPN stimulation” (E). (F) The median values across animals (n=7) have a significant decrease when PPN is stimulated. Illustration in (A) was adapted with permission^54^.

### PPN stimulation and hippocampal theta rhythm

Next, we investigated the effect of PPN stimulation on the hippocampal theta. The theta rhythm was measured using local field potential (LFP) electrodes placed across the first region of the hippocampal circuitry, Cornu Ammonis (CA1), and the dentate gyrus (DG) (**Fig. 2A-B**). Again, the movement was measured using accelerometers, and the PPN-induced arrest in the movement was seen as the acceleration coming to zero (blue-shaded regions, **Fig. 2C**). The hippocampal LFP recording exhibited a strong rhythm before PPN stimulation (**Fig. 2D**), but following the PPN stimulation the rhythm changed. The spectral content changed from having a prominent peak in the theta range (6-9 Hz) to having marginal power in this range (blue band, **Fig. 2E**). This was also reflected in the integrated spectral content in the theta range, which had a diminishing value after the onset of PPN stimulation (**Fig. 2F**). In conclusion, PPN stimulation obstructs the ongoing theta rhythm while also halting the movement.

**Fig. 2.**
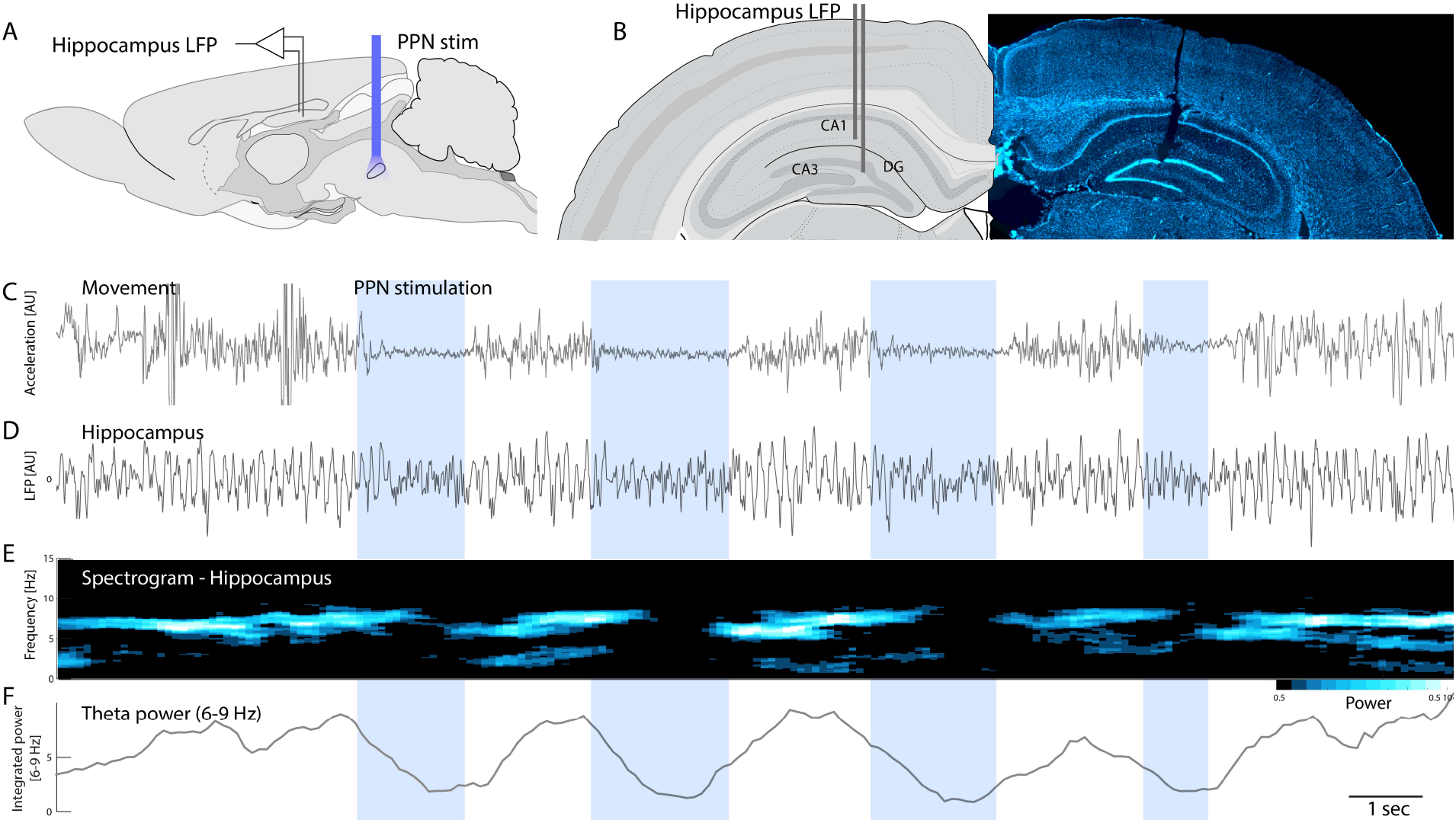
PPN activation obstructs the hippocampal theta rhythm. (A) Sagittal view of the location of the hippocampal LFP electrode and the optical fiber in PPN. (B) Coronal histological section showing the LFP electrode across the dentate gyrus and CA3 (dorsal up, fluorescent DAPI stain). (C) The arrest of movement of the rat is evident by a drop in acceleration following the onset of PPN stimulation (blue-shaded regions). (D) Concurrent LFP activity in the hippocampus with prominent theta rhythm, which evaporates with the PPN stimulation. (E) The spectrogram of the hippocampal LFP displays power in theta range which ceases after PPN stimulation. (F) The integrated theta band power (6-9 Hz). The histological section in (B) is a fluorescent DAPI stain. Illustrations in (A-B) were adapted with permission^54^.

### Decay of the movement arrest and theta rhythm

What is the delay in movement arrest from the onset of PPN stimulation? The time course of the arrest may provide important clues to the neural mechanisms. Hence, we estimated the course of the movement arrests, by fitting an exponential decay, *exp*(*− t/τ*), to the amplitude of the accelerometer data across animals (**Fig. 3A**). The median value of the time constant, *τ*, across events and animals ranged from 90 to 200 ms. One animal (rat 15) had much slower dynamics, which we attribute to the accelerometer not being properly attached to the implant, hence representing a higher upper bound on the time constant. The mean value across animals, when excluding this animal, was 157 ms (horizontal line,**Fig. 3B**). These values are larger but comparable to the values of motor arrest in mice^5^. The larger values of motor arrest are likely due to the larger body weight of rats, which is more than ten times larger than mice. Next, we investigated the timescale of movement arrest together with the cessation of the theta rhythm (**Fig. 3C**). The purpose is to establish a temporal order of the movement arrest compared with the theta rhythm to verify a potential causal relationship. However, due to limitations in the time-frequency resolution, i.e. *the Gabor uncertainty*, the estimation of the change in theta rhythm is difficult to do precisely in time. To mitigate this issue and better compare the temporally well-defined arrest in movement with the less well-defined obstruction of theta rhythm we use sliding windows of equal size in both measures (**Fig. 3D**). We found that both the accelerometer data and the theta rhythm had decay in the activity that was indistinguishable from each other (**Fig. 3D-E**). The standard deviation of the theta rhythm estimation was larger than the movement arrest, again due to the Gabor uncertainty. This is seen as a larger shaded area (cf. cyan and grey **Fig. 3D**) and a wider spread of observations in time constants (cyan vs grey, **Fig. 3E**). Nevertheless, their distributions had a mean, which was not significantly different in 3 out of 4 animals (Wilcoxon signed-rank test, p > 0.05). We conclude that the decay in both theta oscillation and the movement is occurring at a similar rate, which was indistinguishable in our experimental conditions.

**Fig. 3.**
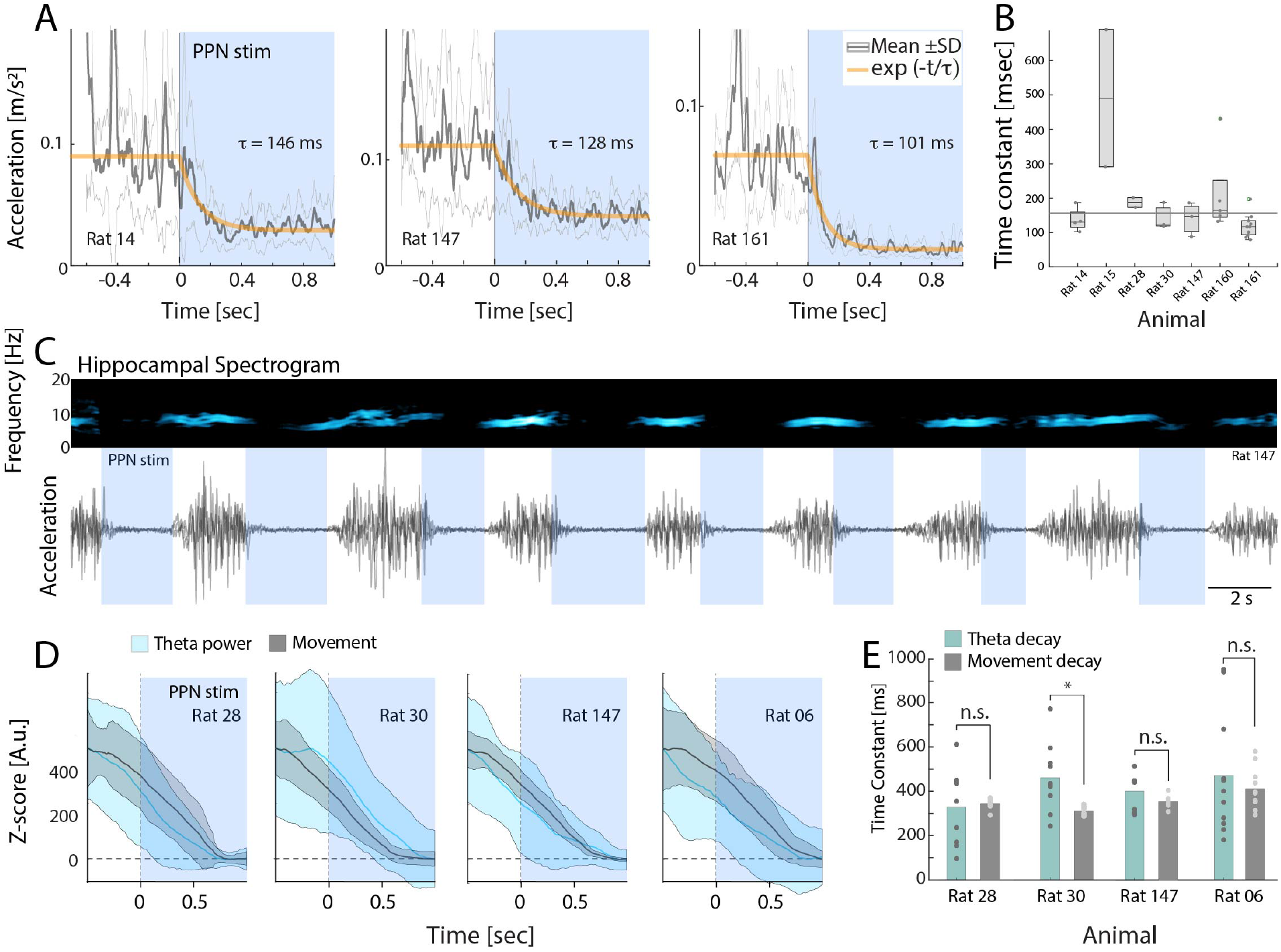
Decay dynamics of movement and theta rhythm following PPN stimulation are similar. (A) The mean movement before and during PPN stimulation (accelerometer measurement ± standard deviation, SD, as the grey lines) for three sample animals. An exponential decay was fitted to the mean (orange line). (B) Whisker plot of time constants of the exponential decay fit for individual instances and animals. The mean time constant was 157 ms across animals (excluding rat 15). (C) A sample trial of hippocampal LFP spectral content (top) during movement and evoked arrest (blue regions), which was measured by accelerometry (bottom). (D) Mean ± SD for the hippocampal theta power (cyan) and accelerometry (grey) for four sample animals before and after the onset of PPN stimulation (blue region). (E) Histogram of time constants of decay of the theta rhythm (cyan) and movement (grey) across animals. There was no significant difference (n.s.) in the mean of time constants for theta rhythm vs. movement in 3 out of 4 animals (Wilcoxon signed-rank test, p>0.05)

### Location of PPN stimulation and cell types

To identify the location of the AAV virus injection, its expression, and optical fiber implantation, histology was performed (**Supplementary Fig. 2 and 3**). Histological brain sections of rats where a successful movement arrest was observed (**Supplementary Fig. 2**) and unsuccessful injections (**Supplementary Fig. 3**). The location of the fiber implantation and AAV virus spread leading to successful movement arrest was verified both using post-mortem MRI-scan and immunohistochemistry (**Fig. 4A-C)**. The PPN is located in the caudal mesencephalon rostrally, dorsally, and ventrally to the decussation of the white matter track coming from the paired superior cerebellar peduncle, which goes towards the thalamus. This white matter track is seen as a slightly darker shadow found at the tip of the vestige from the optical fiber in the structural magnetic resonance imaging (MRI) scan (**Fig. 4B**). The same tissue was sliced and stained using immunohistochemistry and imaged using fluorescent microscopy (**Fig. 4C**). This revealed the area of transfection, where the viral reporter (mScarlet) appears red. Since the PPN is the only nucleus in the area, which contains cholinergic neurons, the staining of cholinergic neurons (ChAT antibody in green) ensures the appropriate location of the fiber and virus. The cholinergic neurons were a few in number, sparsely scattered, and difficult to see, although somata were identified (**Fig. 4D**). To get a better grasp of the location of the part of the PPN, which induces movement arrest, some of the brains were cleared and imaged using light sheet microscopy where the rostral location of the virus injection and optical fiber location are visible with respect to the cholinergic somata (ChAT staining, **Supplementary movie 2**). The location of the successful injection was in the rostral aspects of the PPN nucleus in agreement with previous observations^7^. Next, we investigated what cell types in PPN were transfected and potentially responsible for the behavioral phenotype. Besides cholinergic neurons, the PPN also consists of glutamatergic and GABAergic neurons^15,23,55^. The transfected cells were identified using a combination of immunohistochemistry and RNAscope. The transfected cells that appeared in red (mScarlet) had colocalization with cells, which were both cholinergic and glutamatergic (**Fig. 4E**). Such dual neuronal identity has been reported previously^55,56^. However, most of the transfected cells were glutamatergic as identified with the presence of the gene VGluT2 using RNAscope, which was quantified using Menders correlation (**Fig. 4F**) and manual counting (**Fig. 4G**). Percent of cholinergic neurons transfected was low primarily due to the low number of cholinergic neurons in general. There were also GABAergic neurons among the infected, but the fraction was small (25%). The observation that the majority of infected cells were glutamatergic combined with the induction of movement arrest is in agreement with the observation from mice, where a subset of glutamatergic neurons, which express the gene Chx10, can induce similar movement arrest^4,5^.

**Fig. 4.**
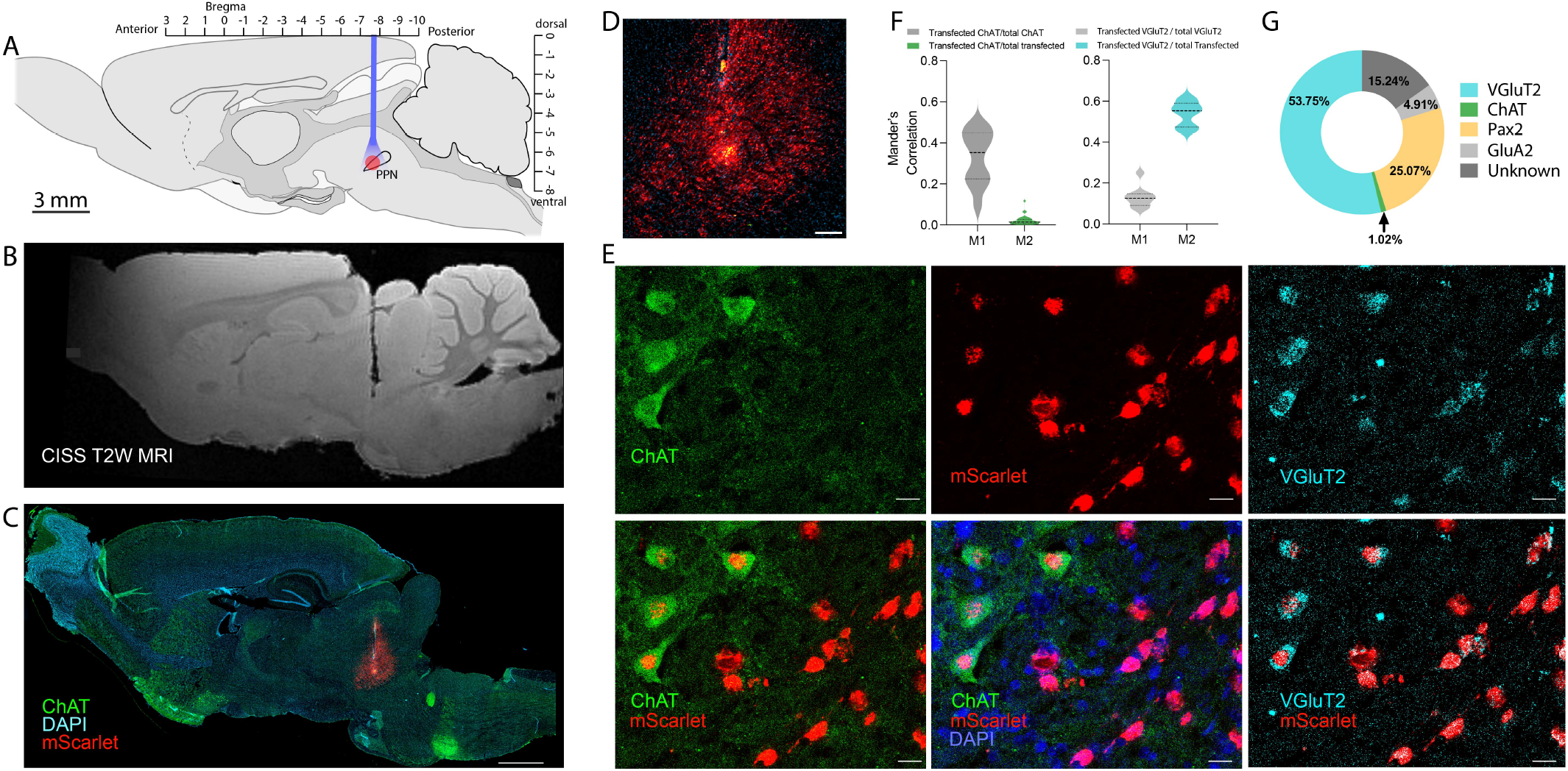
The transfected cells in PPN. (A) The injection of virus and implantation of optical fiber in sagittal stereotaxic coordinates. (B) Post-mortem reconstruction of the location of the optical fiber (MRI-structural scan, track shown in black). (C) histological section of the same tissue as in (B) with the viral reporter (mScarlet using immunohistochemistry) in red, Cholinergic cells (ChAT) in green, and nuclei (DAPI) in blue (scale bar = 1000 *μ*m, cerebellum missing). (D) Zoom-in view of the injection site in (C), shows mScarlet reporter in red (detected using RNAscope) and ChAT cells in green (scale bar = 200 *μ*m). (E) Among the transfected cells were cholinergic cells (ChAT immunohistochemistry, upper and lower left, and middle) and glutamatergic (VGluT2, upper and lower right, in cyan using RNAscope). DAPI was used as a nuclear stain (shown in blue,scale bar = 20 *μ*m).(F) Colocalization was quantified using Manders correlation of a fraction of ChAT (left) overlapping with mScarlet out of all ChAT (M1) and all transfected (M2) and glutamatergic cells (right) overlapping with mScarlet out of all glutamatergic cells (M1) and all transfected cells (M2). (G) Manual counting of colocalization of VGluT2, ChAT, and GluA2 out of all infected cells in percent. N=3 rat brains, n=9-10 sections. Illustrations in (A) were adapted with permission from^54^.

## Discussion

For the brain to perceive events, the location in space has to be matched with the time of a given event^36,37^. The hippocampal formation has a key role in linking the “where” and the “when” during spatial navigation and this is associated with a prominent electrical activity, the theta rhythm. The movement of the body and the commands to contract muscle must be accounted for in the hippocampal formation. Although regions of the brainstem have been known to suppress motor reflexes^57^ and arrest movement^58^, the recent observation that the PPN can induce a global arrest of movement^4,5,7^ introduces a new internal means to tamper with the link between movement and cognition. In this motor hierarchy, how will activation of this lower-level nucleus affect the movement-associated brain rhythm at the hippocampal level? In this report, we tested the impact of PPN-induced movement arrest on the ongoing hippocampal theta rhythm.

The PPN was optogenetically activated to induce movement arrest. The hippocampal theta rhythm was simultaneously recorded and the movement was assessed with accelerometry (**Fig. 1-3**) and videography (**Supplementary Movie 1**). The animal was either roaming around or on a treadmill. During the movement, there was a distinct rhythmic electrical activity in the hippocampus at the 6-9 Hz, which is the theta rhythm. Upon activation of the PPN, this theta rhythm swiftly vanished and the movement was halted. In the attempt to establish a causal order between the halt of execution of movement and the obstruction of theta rhythm, the time constants of these two events were estimated. The decay time of both processes was on the order of 2-300 ms and not distinguishable given our experimental conditions.

How can activation of a low-level brainstem nucleus obstruct the cortical rhythms? The PPN has a subpopulation of cholinergic neurons, which is part of the cholinergic arousal system. The PPN and the arousal system are well-known to desynchronize neocortical activity^39^ and may also be implicated in postural disorders in Parkinson’s disease^59^. Although the cells responsible for the movement arrest are not cholinergic^5,17^, there were indeed cholinergic neurons among the infected cells (**Fig. 4**). Activation of these ascending fibers is also known to induce strong hippocampal theta rhythm, rather than blocking it^39–41,60,61^. For instance, activation of PPN using infused carbachol and electrical stimulation induces hippocampal theta in rats^60,62^. During sleep, we also observed the induction of hippocampal theta rhythm during and following PPN stimulation (data not shown), which is consistent with the previous observation of cholinergic facilitation effect on hippocampal theta rhythm. Hence, the input of ascending fibers from PPN to the hippocampus cannot explain the obstruction of the theta rhythm associated with the arrest of movement. This indicates that the effect on theta rhythm is likely linked to immobility itself.

Immobility is generally associated with the absence of theta rhythm unless the animal is alert during behavioral freezing. If the induced arrest of movement was akin to behavioral freeze the brain should be alert with the associated hippocampus theta rhythm. For this reason, we suggest that the part of the PPN, which can induce movement arrest, represents the motor part of an omnipotent arrest of movement.

## Materials and Methods

### Animals and Ethical statement

Wild-type Long Evans adult male rats (aged 12-24 weeks, 450-600 g) were used to perform the experiments. The rats were obtained from Charles River Laboratories and housed in cages equipped with bedding, food, water, temperature and air sensors and kept in 12h light/ 12 h dark cycle. Before any surgical procedure, the rats were housed in pairs for one week in the animal housing facility for acclimatization. All the experimental procedures were performed in compliance with with ARRIVE guidelines and the Council of the European Union (86/609/EEC) and authorized by the Danish veterinary and food administration (animal research permission number 2019-15-0201-00018).

### Surgery, virus injection, and fiber implantation

All the surgical procedures on Wild-type adult rats were performed in aseptic conditions. The rats were anesthetized using gas anesthesia (isoflurane in a mixture of air and oxygen). The surgery room was divided into 2 sections: a dirty and a clean section. In the dirty section, an initial part of the procedure was performed such as shaving hair off the rat’s surgical area and cleaning the shaved area with chlorhexidine (from inside to outside) followed by cleaning with ethanol (70%). Later, the rat was moved to the clean section, which is the surgery table with a stereotaxic robot, on a heating pad with a temperature sensor, and an oximeter was used to monitor the oxygen level during the surgery. The head of the rat was fixed using the ear bars. Ocryl gel was used to keep the eyes moist and the surgical area was again cleaned with 70% ethanol. Then a sterile drape was placed on the anesthetized rat and a hole was made on the drape at the surgical area of interest and Mepitel film with a fine cut in the middle (Molnlycke Healthcare, Goteborg, Sweden) was placed on the shaved sterile area of the rat.

An incision was made on the skin and bulldog forceps were used to stretch the skin and increase the visibility of the skull. The skull was made completely dry using electrocautery and hydrogen peroxide. The skull was painted with a thin layer of dental cement (C&B Metabond quick adhesive cement system, Parkell). A 3D-printed head plate with an attached copper mesh^**?**^ was glued on the skull, craniotomy was performed and an AAV virus (AAV9-CamKIIa-ChrimsonR-mScarlet-Kv2.1, which was a gift from Christopher Harvey via Addgene.org by^63^ was injected with a glass capillary using a stereotaxic robot (Neurostar GmbH, Tubingen, Germany) into the PPN (500 nl injection volume, Fig. 1A, coordinates (DV: 7.8, ML: 1.58, AP: -7.0). An optical fiber was subsequently implanted 200 micro-m above the injection site and the hole was closed using cyanoacrylate glue followed by dental cement. A copper crown was built out of the mesh using a thin layer of dental cement to protect the protruding optical fiber ferrule. The crown also served as a Faraday cage during electrical recordings. The post-operative care was given where the rats were given buprenorphine mixed with Nutella^64^(sublingual tablets 0.2 mg crushed to powder and mixed with 1 g Nutella, dose = 0.4 mg/kg, every 12 h for 3 days) and carprofen (subcutaneously (s.c.), 5 mg/kg, once a day for 5 days) as analgesic and anti-inflammatory and Baytril (s.c., 5 mg/kg, once/day for 10 days) as antibiotic drugs. The rat was monitored twice/day for 3 consecutive days followed by once/day for 7 days.

### Measurement of theta rhythm: Implantation of electrocorticographic electrodes

3-4 weeks after the injection of the AAV virus, the virus shows expression. Therefore, after 4 weeks the implant was tested for whether it could induce movement arrest when using an optogenetic pulse train delivered to PPN with a red LED laser (625 nm, Model M625F2, Thorlabs, Inc.). After confirming the motor arrest, a second surgery was performed where 4 tungsten electrodes were implanted in the primary motor cortex (AP =1.56-2.52, ML =2.2-2.3 and DV=2.0-2.3) and 4 in the hippocampus (AP = -3.3-3.6, ML = 2.0 and DV= 3.0-3.4) together with 1 reference silver and 1 ground silver electrodes. Post-surgical care was given as reported by^64^.

The Teflon-coated tungsten or silver wires were inserted into the hippocampal formation in the CA1-CA3 where the local field potential is strongest^65,66^. The electrical signals were recorded using a miniaturized head stage with integrated accelerometers tethered with a lightweight interface cable to the amplifier (RHD 32 channel head stage, Intan technologies). The movement was measured using accelerometers available on the head stage. The reading, which was in Volt was converted to acceleration as 340 mV/g, where 1g is 1 m/s^2.

### RNAscope

At the end of the experiment, the rats were perfused with 4% formaldehyde. The perfusion procedure was adapted from previous studies^64^. The brains were extracted and fixed again using formaldehyde for 4 h followed by cryoprotection using 30% (w/v) sucrose for 48 h. Later the brains were stored at -80°C. 20 *μ*m thin brain sections were collected on Superfrost plus glass slides (Thermo Fisher Scientific GmbH, Germany) and stored at -20°C until further use. In situ labeling of targeted mRNA was done by RNAscope method, using a commercially available kit (Advanced Cell Diagnostics, Hayward, CA) according to the manufacturer’s protocol. Dehydrated tissue sections were blocked with hydrogen peroxide and boiled in target retrieval reagent for 5 min. Then the tissue was incubated with protease plus for 30 min at 40C and targeted probes against mScarlet (Cat No. 572421) and VGluT2 (Cat No. 319171-C3) (Advanced Cell Diagnostics, Hayward, CA) were hybridized on the tissue for 2 h in Hybez oven (Advanced Cell Diagnostics, Hayward, CA) followed by a series of signal amplification and washing steps. The signal was visualized by incubation slides with OpalTM570 and OpalTM690 reagents (1:1000) (Perkin Elmer). Then sections were processed for immunohistochemistry.

### Immunohistochemistry

After completing the RNAscope procedure, the same brain sections were used to perform immunohistochemistry. This procedure was adapted from prior studies^67,68^. The slides with brain sections were washed in 1X phosphate-buffered saline (PBS) followed by incubated them with Blocking solution (5% bovine serum albumin, 5% fetal bovineserum, 0.3% Triton X-100, 1% PBS) for 2 hours at room temperature followed by administration of primary antibodies such as ChAT (goat polyclonal, 1:500, EMD Millipore AB144P), Pax2 (rabbit polyclonal, 1:500 dilution, Invitrogen UD283859) and GluA2 (mouse monoclonal, 1:200 dilution, EMD Millipore MAB397) for overnight at 4°C. The next day, the slices were washed and incubated with the following secondary antibodies (1:500): donkey anti-goat Alexa Fluor 647 (Abcam, AB150135), donkey anti-rabbit Alexa Fluor 647 (Invitrogen, A31573), donkey anti-rabbit Alexa Fluor 488 (Invitrogen, A11055), donkey anti-mouse Alexa Fluor 647 (Invitrogen, A31571) and 4’,6-diamidino-2-phenylindole (DAPI, 1:1000; Sigma-Aldrich) at room temperature for 2 hours. DAKO mounting medium was used to mount the slides followed by visualizing under Axio scan Z1 and confocal microscope using 20X magnification. The images captured by microscopes were analyzed using Zen Lite 3.1 and ImageJ software. The correlation coefficient was computed by using the Manders correlation coefficient and metric matrix^69^. Adobe Illustrator was used to create a contour of the brain section/s to show the viral injection site (**Fig. 1A, Supplementary Fig. 2-5**). Graph-pad prism was used to create graphs.

### Tissue clearing and light sheet microscopy

To view the viral expression in 3D one rat brain was cut in half and cleared using a 2nd generation ethyl-cinnamate clearing method (2ECi clearing, **Supplementary Movie 2**)^70^ and the other brain was cut from the sides sagittally and cleared using Adipo-clear method (**Supplementary Movie 2**)^71^. During the 2ECi procedure, the brain was stained with a nucleic acid stain (Sytox, green nucleic acid stain, 1:5000, ThermoFisher Scientific, cat. S7020). The brain was imaged on an LCS Spim light-sheet microscope (Bruker). The endogenous fluorescent reporter (mScarlet^72^) was visible as a red fluorophore (**Supplementary Movie 2**) During the Adipo-clear procedure, the brain was stained with primary antibodies, ChAT (goat polyclonal, 1:400, EMD Millipore AB144P) and RFP (rabbit, 1:1000, Rockland) followed by incubation with secondary antibodies, donkey anti-goat (1:500, Alexa fluor 647) and donkey anti-rabbit (1:500, Alexa fluor 555). The cleared brain was imaged with a Zeiss LS7 microscope (**Supplementary Movie 2**).

### Magnetic Resonance Imaging

Structural Magnetic resonance imaging (MRI) was done for post-mortem anatomical reconstruction and verification of electrode and optical fiber location. After transcardial perfusion using PBS followed by paraformaldehyde (PFA, 4%) the tissue was kept overnight in PFA followed by transferring to sucrose solution (30% w/v) and finally to PBS. Later the brain was transferred to a tube containing fluorinert (FC-40, Sigma-Aldrich) to reduce the background signal. The tube was placed in a 9.4 T pre-clinical MRI scanner (BioSpec 94/30; Bruker Biospin, Ettlingen, Germany) equipped with a 1.5 T/m gradient coil. A T2-weighted scan sequence was performed, the Constructive Interference in Steady State (CISS) in isotropic resolution of 50 um (**Fig. 4B** and **Supplementary Movie 2**).

### Data analysis

The spectrogram was calculated using the multi-taper method described in the previous report^66^. The LFP potential recordings from the hippocampus were filtered using custom-designed Matlab code (Mathworks 2020b) consisting of a 3-pole Butterworth filter. The time constant for the decay in the movement was estimated from the full-wave rectified accelerometer measurement, which was averaged across x, y, and z directions and smoothed. The decay was fitted to an exponential decay model, exp(-t/*τ*), where *τ* presents the decay time constant.

## Acknowledgements

Funding: This work was supported by The Independent research fund Denmark, and the Carlsberg foundation. Thanks to Yuki Mori from Center for translational medicine for performing the MRI scans. Thanks to Palle Koch and Jakob F. Sørensen for help designing and constructing the treadmill.

## Author contributions

R.W.B. and J.K. conceived and designed the experiments. J.K. performed the surgeries and data analysis of post-mortem tissue. J.K., R.W.B. performed the experiments. J.K. and M.B. performed immunohistochemistry. O.D. performed RNAscope.

J.K. and G.H. performed tissue clearing. R.W.B. and S.K. analyzed the electrophysiology data. R.W.B. wrote the manuscript.

