## Supplementary information for "Arrest of movement induced by Pedunculopontine-stimulation obstructs hippocampal theta rhythm"

Jaspreet Kaur[1], Salif Komi[1], Oksana Dmytriyeva[1], Grace A. Houser[1], Madelaine C. A. Bonfils [1], and Rune W. Berg[1]

[1]Affiliation not available

August 25, 2023

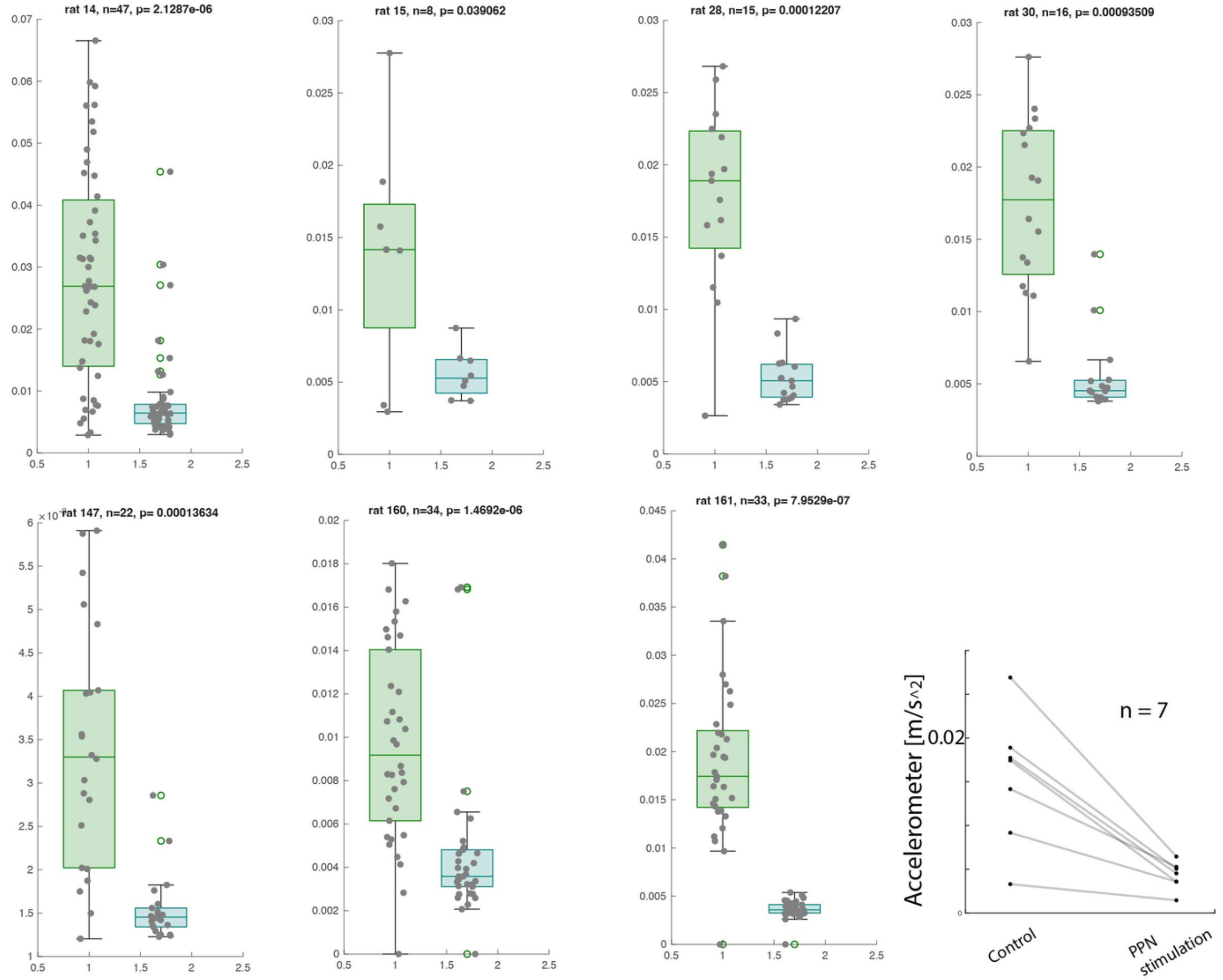


Figure 1: **Accelerometer recordings while locomotion versus movement arrest:** Accelerometer recordings obtained from 7 rats where the green box plot show (in each plot) acceleration when the rat was locomoting whereas the blue box plot shows nullified acceleration while the optogenetic stimulaiton of PPN is on. The line plot shows accelerometer recordings when the rats were locomoting (as control) versus the movement arrest (PPN stimulation).

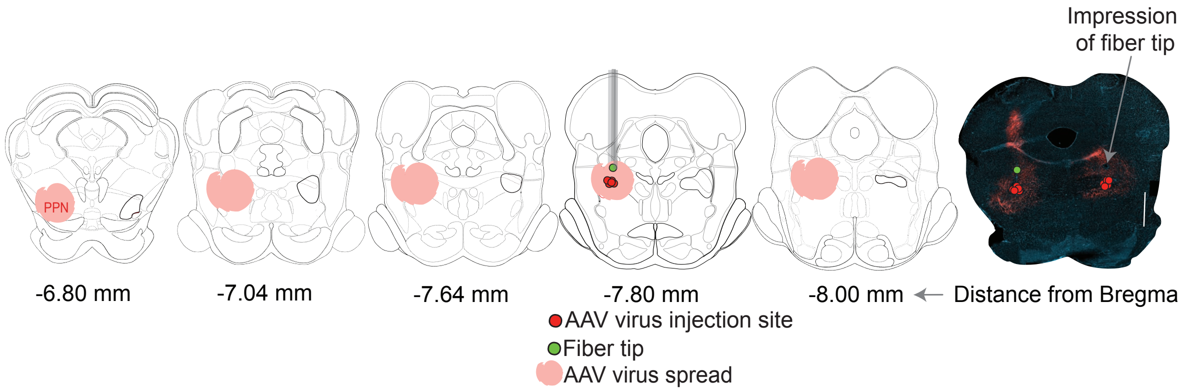


Figure 2: **Coronal brain sections highlighting the AAV virus injection site and optical fiber implant site in PPN.** Coronal brain sections highlight the PPN location in the various brain sections. AAV virus was injected at a -7.8 mm distance from bregma (shown with red circles), the virus spread has shown in pink and the optic fiber implant site has illustrated in green circle (adapted from (Swanson, 2018) with permission). The right image shows the coronal histology section of the brain with an AAV virus injection (red circles), AAV virus spread (in mScarlet in red) and impression of fiber tip (with an arrow and green circle).

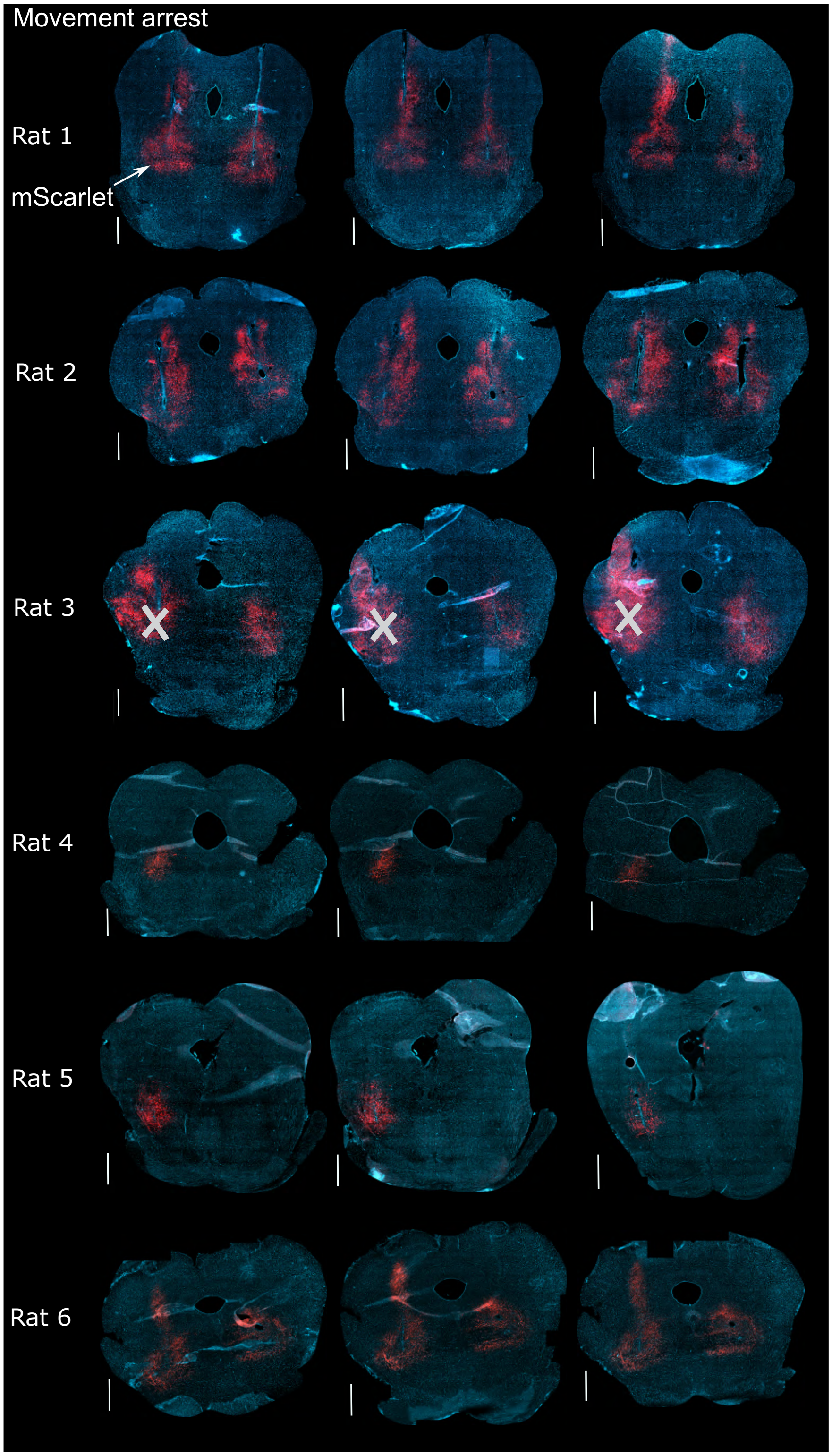


Figure 3: **Histology sections of brains with movement arrest.** Coronal brain sections from 6 rats (3 sections per rat) showed clear movement arrest when PPN (where a mScarlet protein, in red, from AAV virus was expressed) was optically stimulated. Stimulation in Rat 3 only elicited in movement arrest when stimulating on one side, which was the side opposite to where the white cross is. The side with the white cross was unresponsive. Scalebar = 20 µm.

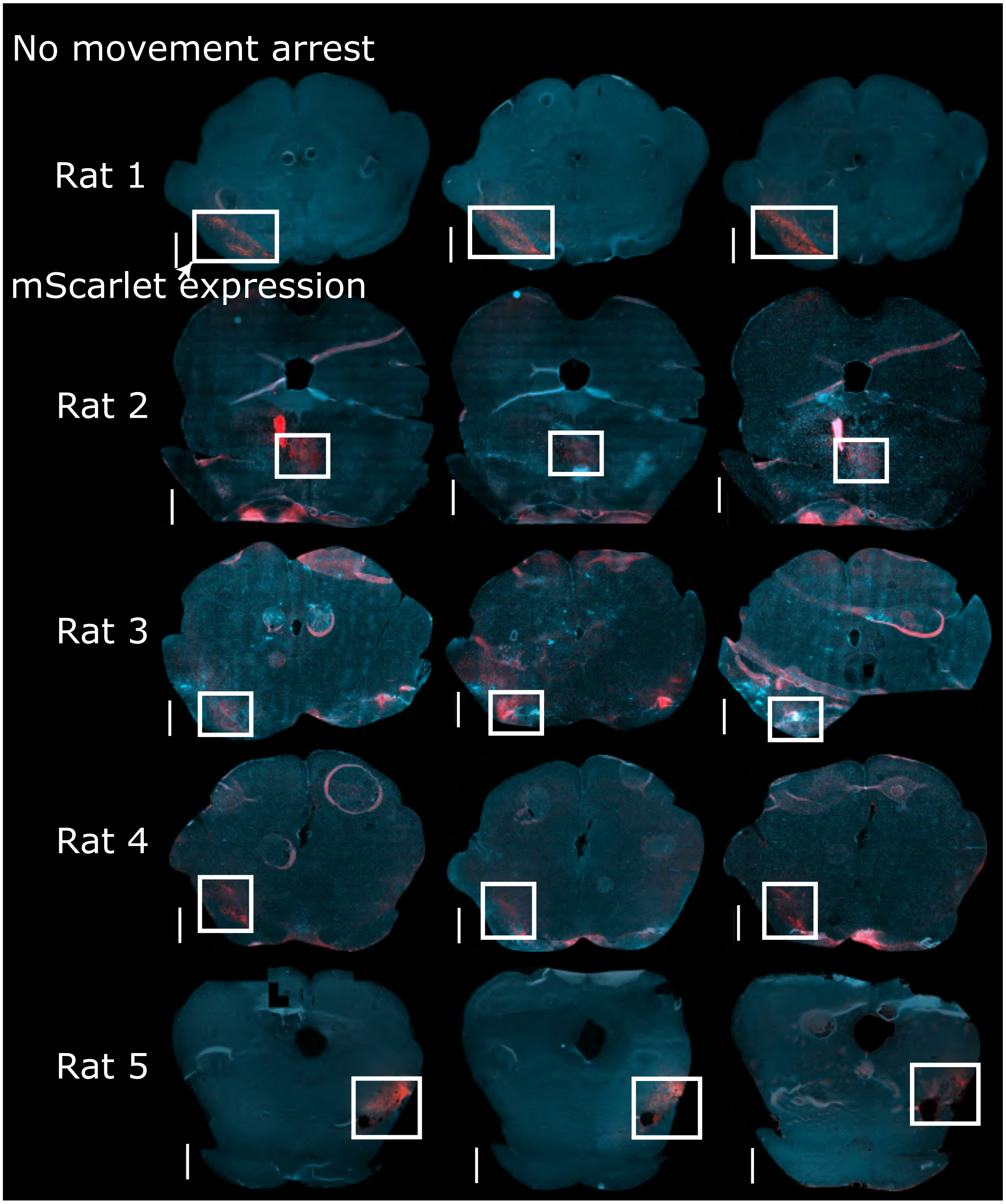


Figure 4: **Animals with AAV injections, which did not induce movement arrest.** Each row show three slices from one animal where the mScarlet from AAV was expressed (white boxes highlight mScarlet expression) and the optical fiber with implanted, but the optogenetic stimulation did not lead to movement arrest or any other overt response.

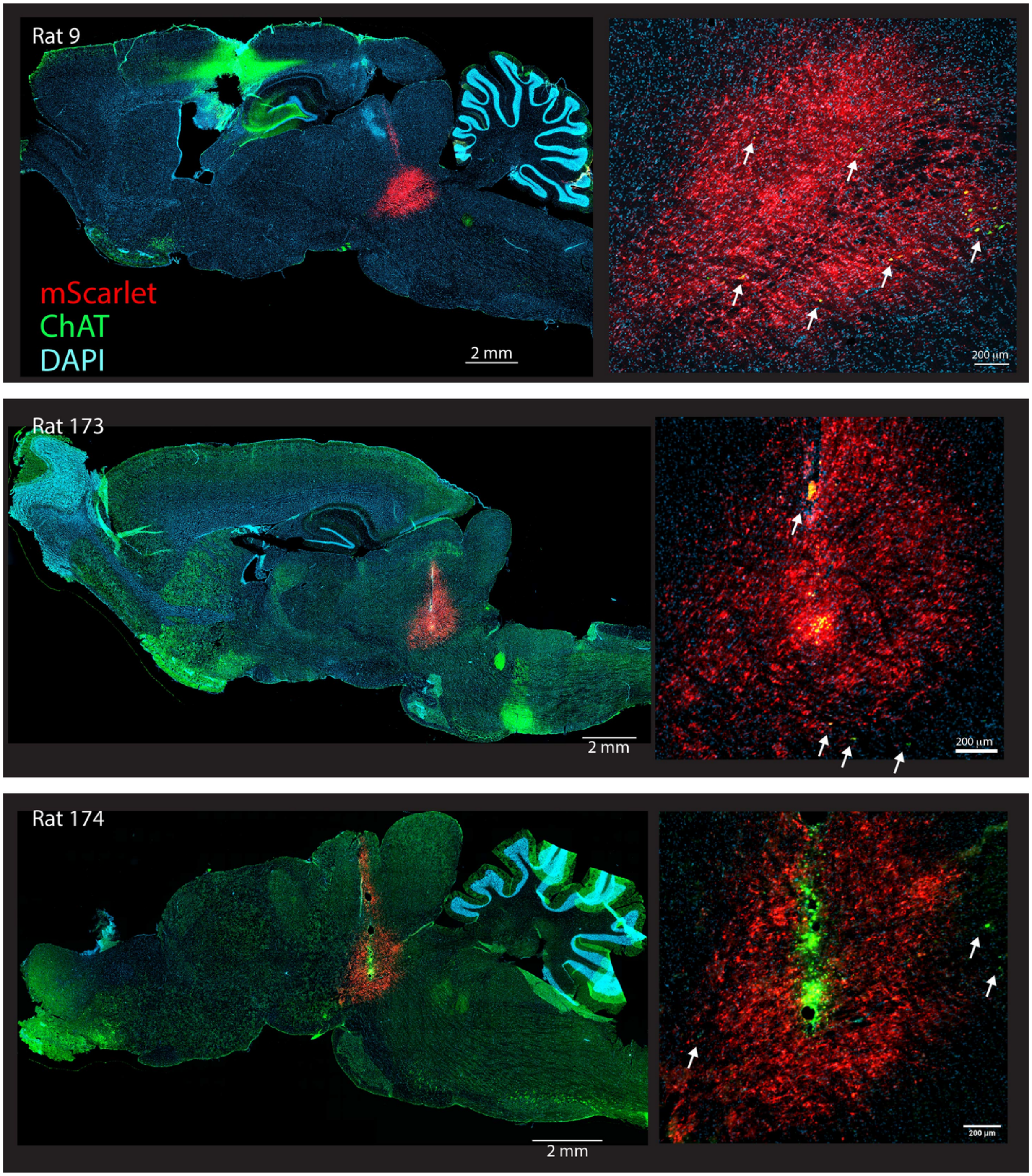


Figure 5: **Location of PPN, viral injection, and optical fiber in sagittal section.** Left: Immunohistochemical sections (thickness: 20 µm) for 3 animals, showing the viral reporter (mScarlet) in red, the cholinergic neurons (ChAT in green) and neuronal nuclei (DAPI, cyan). Right: enlargement of the infected region. Scattered cholinergic somata (white arrows) confirm the overlap with PPN. Images have been contrast-enhanced for better visual illustration. Animals: Rat 9, 173 and 174.

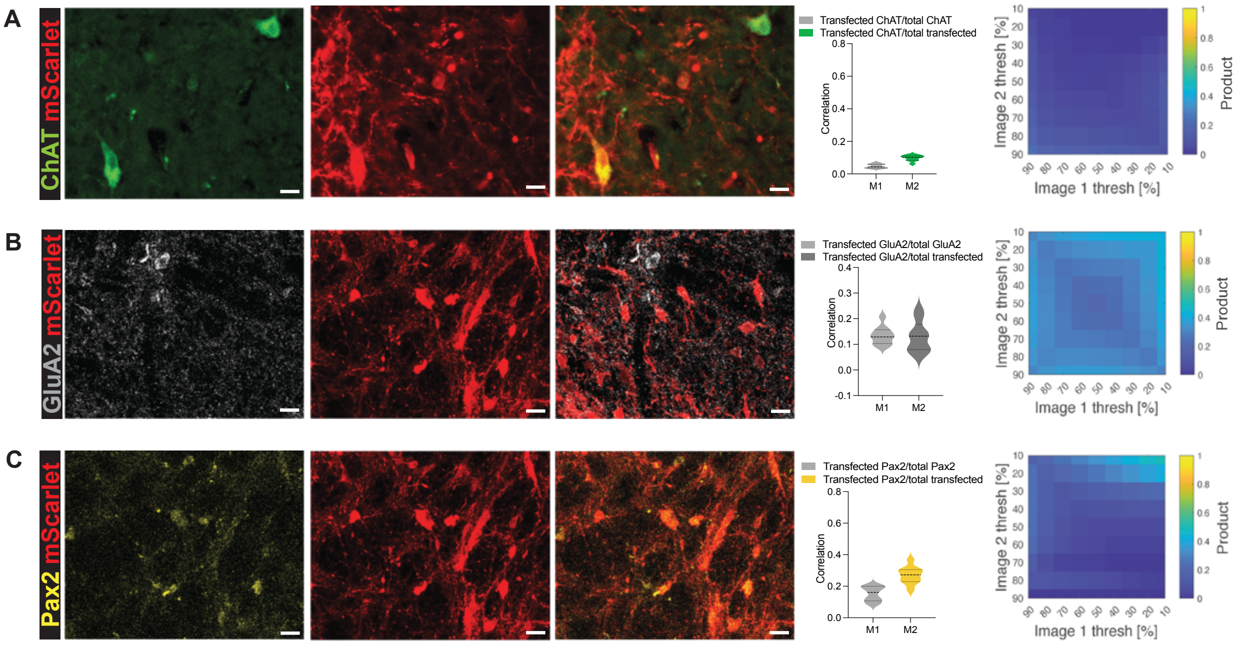


Figure 6: **Low colocalization of mScarlet with ChAT, GluA2, and Pax2.** A-C) Left panels: Immunohistochemistry images showing low co-expression of mScarlet protein with ChAT, GluA2, and Pax2 biomarkers. Middle: Violin plots of Manders correlation (M1 and M2) values for the three biomarkers with mScarlet. Right: Heat map illustrating correlation at different thresholds where the heat map for ChAT (A) showed lowest, for GluA2 (A) depicted low, and for Pax2 (C) represented medium correlation.

**Supplementary Movie 1. PPN stimulation arrests movement and obstructs hippocampal theta rhythm.** During treadmill locomotion by a rat, the hippocampal local field potential has a prominent spectral content in the theta band (blue band shown in the spectrogram, middle panel). The movement is also measured using accelerometry (bottom panel). The PPN is periodically stimulated with light, to activate an expressed opsin (ChrimsonR). The stimulation is indicated by a blue light appearing at the lower right, top panel, and gray-shaded regions in the accelerometry measurement. Animal: Rat 147.

**Supplementary Movie 2. Location of the PPN, virus injection, and optical fibers.** The movie has three parts: 1) Expression of virus in a cleared half-brain. The rat brain was cleared using ECi clearing and the cell nuclei were stained using sytox green staining (cyan). The tissue was imaged on a light sheet microscope (LCS SPIM Bruker). Animal: Rat 93. 2) A magnetic resonance imaging scan of the post-mortem rat brain both in sagittal and coronal plane, shows the tracks of where the optical fibers were located. The PPN is located around the white fiber bundle (seen as a darker grey shadow) coming from the superior cerebellar peduncle. Animal: Rat 173. 3) Tissue-cleared rat brain using Adipo-Clear, which is also immunostained using ChAT- (green) and mScarlet-(red) antibodies. This shows both the location of the virus injection (in red), the location of PPN (green ChAT staining), and the visible tracks from autofluorescence from where the optical fibers were located. Animal: Rat 12.
